# Structure, activity and inhibition of human TMPRSS2, a protease implicated in SARS-CoV-2 activation

**DOI:** 10.1101/2021.06.23.449282

**Authors:** Bryan J. Fraser, Serap Beldar, Almagul Seitova, Ashley Hutchinson, Dhiraj Mannar, Yanjun Li, Daniel Kwon, Ruiyan Tan, Ryan P. Wilson, Karoline Leopold, Sriram Subramaniam, Levon Halabelian, Cheryl H. Arrowsmith, François Bénard

## Abstract

Transmembrane protease, serine 2 (TMPRSS2) has been identified as key host cell factor for viral entry and pathogenesis of SARS-coronavirus-2 (SARS-CoV-2). Specifically, TMPRSS2 proteolytically processes the SARS-CoV-2 Spike (S) Protein, enabling virus-host membrane fusion and infection of the lungs. We present here an efficient recombinant production strategy for enzymatically active TMPRSS2 ectodomain enabling enzymatic characterization, and the 1.95 Å X-ray crystal structure. To stabilize the enzyme for co-crystallization, we pre-treated TMPRSS2 with the synthetic protease inhibitor nafamosat to form a stable but slowly reversible (15 hour half-life) phenylguanidino acyl-enzyme complex. Our study provides a structural basis for the potent but non-specific inhibition by nafamostat and identifies distinguishing features of the TMPRSS2 substrate binding pocket that will guide future generations of inhibitors to improve selectivity. TMPRSS2 cleaved recombinant SARS-CoV-2 S protein ectodomain at the canonical S1/S2 cleavage site and at least two additional minor sites previously uncharacterized. We established enzymatic activity and inhibition assays that enabled ranking of clinical protease inhibitors with half-maximal inhibitory concentrations ranging from 1.7 nM to 120 μM and determination of inhibitor mechanisms of action. These results provide a body of data and reagents to support future drug development efforts to selectively inhibit TMPRSS2 and other type 2 transmembrane serine proteases involved in viral glycoprotein processing, in order to combat current and future viral threats.

**SUMMARY PARAGRAPH:** Viruses hijack the biochemical activity of host proteins for viral invasion and replication. Transmembrane protease, serine-2 (TMPRSS2) is a surface-expressed protease implicated in the activation of influenza A, influenza B, and coronaviruses, including SARS-CoV-2, to drive efficient infection of the lungs^1–5^. TMPRSS2 is an attractive target for antiviral therapies, as inhibiting its proteolytic activity blocks efficient viral entry^5,6^. However, a structural and biochemical understanding of the protease has remained elusive and no selective inhibitors are available. We engineered on-demand activatable TMPRSS2 ectodomain and determined the 1.95 Å X-ray crystal structure of the stabilized acyl-enzyme after treatment with nafamostat, a protease inhibitor under investigation as a COVID-19 therapeutic. The structure reveals unique features of the TMPRSS2 substrate recognition pocket and domain architecture, and explains the potent, but nonselective inhibition by nafamostat. TMPRSS2 efficiently cleaved the SARS-CoV-2 S protein at the canonical S1/S2 site as well as two minor sites previously uncharacterized. We further established a robust enzymatic assay system and characterized inhibition by two additional clinical protease inhibitors under study for COVID-19, camostat and bromhexine. Our results provide a body of data and reagents to enable ongoing drug development efforts to selectively inhibit TMPRSS2 and other TTSPs involved in viral glycoprotein processing, in order to combat current and future viral threats.

## MAIN

### Production and structure of on-demand activatable TMPRSS2 ectodomain

TMPRSS2 is a type 2 transmembrane serine protease (TTSP) comprised of an intracellular domain, single-pass transmembrane domain, and a biologically active ectodomain with three subdomains: a low density lipoprotein receptor type-A (LDLR-A) domain, a Class A Scavenger Receptor Cysteine-Rich (SRCR) domain and a C-terminal trypsin-like serine peptidase (SP) domain with a canonical Ser441-His296-Asp345 catalytic triad (Fig. 1a and e)^7,8^. As TMPRSS2 is synthesized as a single-chain proenzyme, or zymogen, it requires cleavage at a conserved Arg255-Ile256 peptide bond within its SRQSR255↓IVGGE activation motif (cleavage site denoted with an arrow) to achieve full maturation of its enzymatic activity^8,9^. We achieved this in high yield by replacing SRQSR255↓ with an enteropeptidase-cleavable DDDDK255↓ sequence to prohibit auto-activation, allowing purification of a secreted form of the full TMPRSS2 ectodomain zymogen from insect cells, analogous to a strategy used for the TTSP, matriptase (Methods)^10^. Subsequent proteolytic activation with recombinant enteropeptidase afforded highly active, homogenous TMPRSS2 to milligram yields and was accordingly named directed activation strategy TMPRSS2 (dasTMPRSS2; Fig. 1b; Extended Data Fig. 1). We determined the X-ray crystal structure of dasTMPRSS2 refined to 1.95Å resolution after acylation of the catalytic Ser441 residue with nafamostat, a broad-spectrum synthetic serine protease inhibitor^11^. We obtained clear electron density for residues 149-491 spanning the SRCR and SP domains but not residues 109-148 containing the flexible LDLR-A domain responsible for linking the protease to the plasma membrane (Fig. 1c). The engineered DDDDK255 activation motif was not resolved in the structure but rather terminated in an unstructured loop, consistent with matured TMPRSS13 (6KD5) and hepsin (1Z8G) structures containing their native activation motifs^12,13^. The newly exposed N-terminal Ile256 of the SP domain formed a salt bridge with the side chain of Asp440, confirming full maturation of the activation pocket, and taken together with the Cys244-Cys365 interdomain disulfide, confirms that this structure represents the bioactive, stabilized form of the protease (Fig. 1d-e).

**Figure 1.**
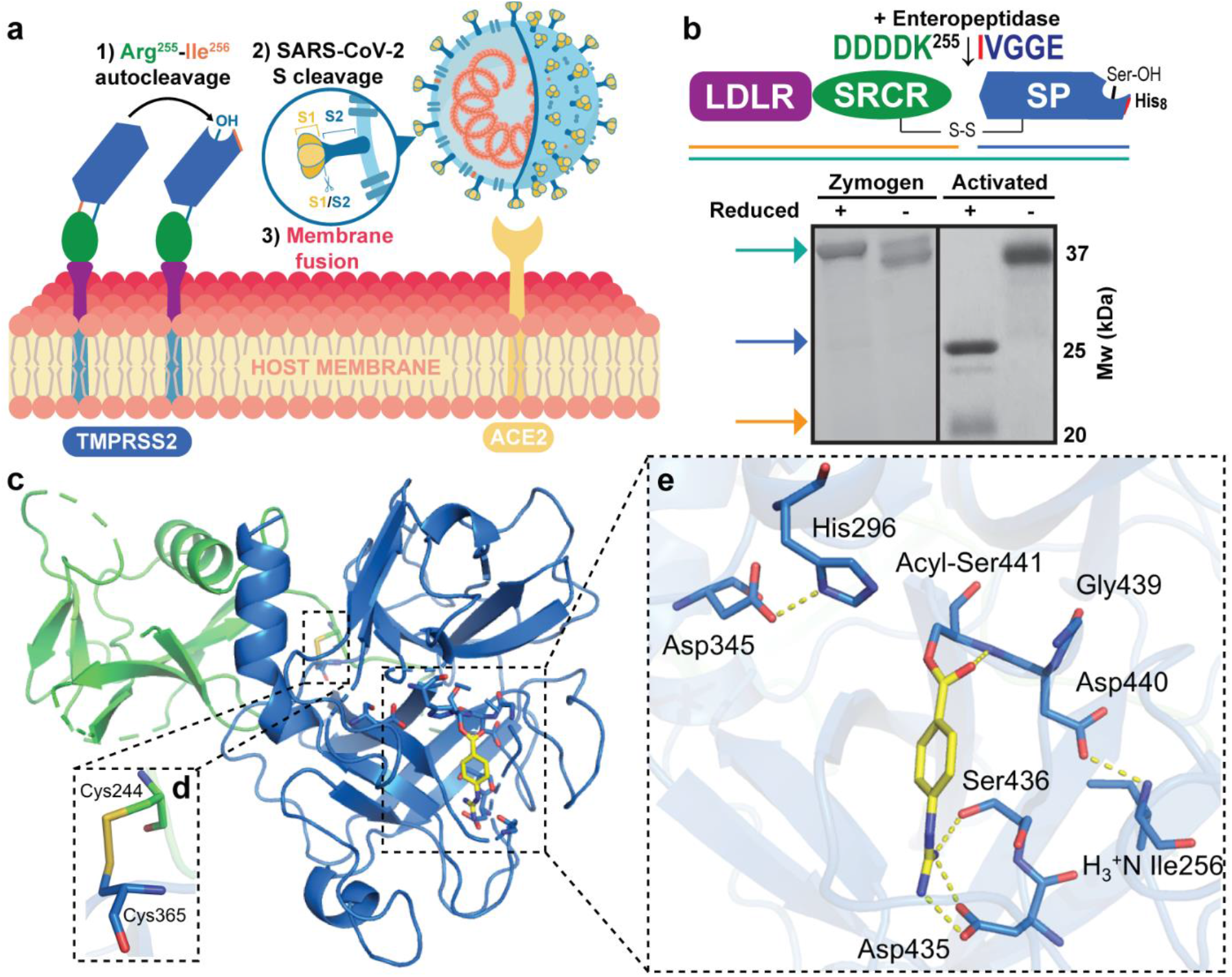
Engineered activation and structural characterization of stabilized TMPRSS2 ectodomain. **a** Full-length, membrane bound TMPRSS2 zymogen undergoes autocleavage activation at the Arg255-Ile256 peptide bond and the matured enzyme proteolytically processes SARS-CoV-2 Spike protein docked to the ACE2 receptor to drive viral membrane fusion. **b** Engineered recombinant TMPRSS2 ectodomain containing the low-density lipoprotein receptor type-A (LDLR) domain, a Class A Scavenger Receptor Cysteine-Rich (SRCR) domain and a C-terminal trypsin-like serine peptidase (SP) domain, features an enteropeptidase-cleavable DDDDK^255^ substitution to facilitate controlled zymogen activation. The non-catalytic (LDLR+SRCR) and catalytic (SP) chains are tethered by a disulfide bond and the activation status can be interrogated by SDS-PAGE under non-reducing and reducing (5% β-mercaptoethanol) conditions. **c** X-ray crystal structure of activated TMPRSS2 ectodomain pre-treated with nafamostat (yellow sticks). **d** The interdomain disulfide pair (Cys244-Cys365) maintains covalent attachment of the SRCR and SP domains. **e** Close-up view of the SP catalytic triad residues (His296, Asp345 and Ser441) and the post-activation Asp440:Ile256 salt bridge showing complete maturation of the protease. Nafamostat treatment results in phenylguanidino acylation of Ser441. Polar contacts are shown as yellow dashed lines.

### TMPRSS2 has a unique and accommodating substrate binding cleft

The TMPRSS2 SP domain is highly conserved with all TTSPs and conforms to the canonical chymotrypsin/trypsin fold with two six-stranded beta barrels converging to a central active site cleft harboring the catalytic triad (Fig. 1c)^14^. Divergent protein substrate specificity of these closely related proteases is conferred through highly variable, surface-exposed loops, denoted Loop A-E and Loops 1-3 (Extended Data Fig. 2)^14^. Unique subsites formed on the face of the SP domain, S4-S3-S2-S1-S1’-S2’-S3’-S4’ recognize substrate P4-P3-P2-P1↓P1’-P2’-P3’-P4’ amino acid positions spanning the scissile bond (Fig 2a; Extended Data Fig 3a). To rationally assign these subsites for TMPRSS2, we superposed the peptide-bound hepsin and TMPRSS13 SP domains (40.1% and 41.4% sequence identity of their SP domains, respectively) belonging to the same hepsin/TMPRSS subfamily as TMPRSS2. The S1 position of TMPRSS2 is occupied by the phenylguanidino moiety of nafamostat, forming salt bridges with the highly conserved Asp435, Ser436, and Gly464 residues in the same binding mode as the guanidino of P1 Arg residues observed in hepsin and TMPRSS13 (Fig. 1e; Fig. 2a; Extended Data Fig. 3a). The TMPRSS2 S2 subsite has a distinguishing Lys342 residue that likely confers a preference for small and/or electronegative P2 substrates, similar to the S2 Lys in enteropeptidase which prefers P2 Asp residues^15^. The S3 and S4 subsites appear open to accommodate various P3 and P4 amino acids and may make favorable receptor contacts with the respective Gln438 and Thr341 positions (Fig. 2a, Extended Data Fig. 3a). On the N-terminal side of the scissile bond, the buried S1’ site appears to accept small, hydrophobic P1’ residues. Overall, the TMPRSS2 active site appears capable of binding various substrate sequences with the strictest preference for the P1 and P2 positions.

**Figure 2.**
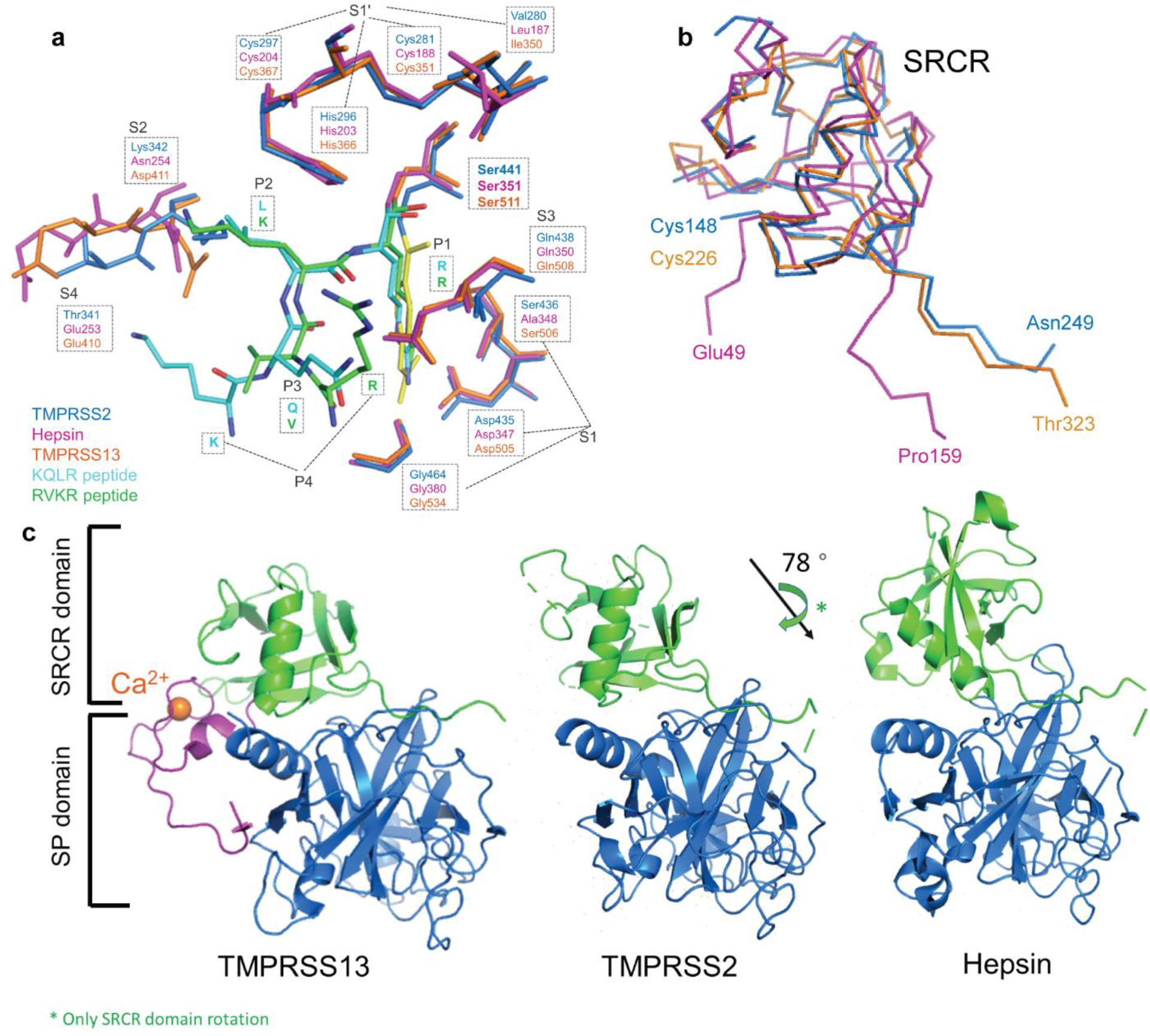
Divergent surface properties of TMPRSS2 inform putative substrate preferences and relative domain organization. **a** The subsites of TMPRSS2 (blue) superimposed on the corresponding residues of Hepsin (magenta, PDB: 1Z8G) and TMPRSS13 (orange, PDB: 6KD5). The S1 subsite occupied by the phenylguanidino acyl group has well conserved Ser441, Asp435, and Gly464 residues, whereas discriminatory residues in S2 (Lys342) and S4 (Thr341) are not occupied. **b** Ribbon representation of the superimposed SRCR domains of TMPRSS2 (blue), Hepsin (magenta, PDB: 1Z8G) and TMPRSS13 (orange, PDB: 6KD5). **c** Relative orientation of the SRCR (green) and serine protease domains (blue) of TMPRSS2, Hepsin (PDB: 1Z8G), and TMPRSS13 (PDB: 6KD5). The LDLR domain of TMPRSS13 is shown in magenta with bound calcium in orange.

Among TTSPs, the SP domain of TMPRSS2 uniquely possesses 3 disulfides and a single unpaired cysteine residue, Cys379 (Extended Data Fig. 2b-d). In all other TTSPs this position forms a disulfide bond with an additional Cys at the equivalent of Thr447, or both cysteines are absent. This unpaired cysteine is conserved in feline, bovine, mouse, and rat TMPRSS2 orthologs (Extended Data Fig. 2e). Furthermore, the unpaired Cys379 is bordered by an expansive 360 Å^2^ patch of exposed hydrophobic surface area in our structure that may serve as an interaction hub for TMPRSS2 binding partners (Extended Data Fig. 2b).

### The SRCR domain confers additional diversity for molecular recognition

The SRCR domain is found enriched in proteins expressed at the surface of immune cells as well as in secreted proteins, and are thought to participate in protein-protein interactions and substrate recognition^16^. The Class A SRCR domain of TMPRSS2 is located on the backside of the SP domain away from the active site and is structurally similar to that of TMPRSS13 despite sharing only 19% sequence identity (Fig. 2b). These two SRCR domains adopt a compact, globular fold with similar orientations relative to their SP domains (Fig. 2b,c). The SRCR of hepsin (7.5% sequence identity) diverges significantly from TMPRSS2/13 with three intra-domain disulfides and a tighter SRCR:SP association dominated by complementary electrostatic patches and buried surface area (Fig. 2c). These conformational differences may play a role in the orientation of the SP domain relative to the plasma membrane as well as modulate activity through recognition or recruitment of partner proteins.

### TMPRSS2 displays robust in vitro peptidase activity

To evaluate TMPRSS2 inhibitors and provide groundwork for future structure activity relationship (SAR) studies, we established in vitro proteolytic activity and inhibition assays. The generic TTSP fluorogenic peptide substrate Boc-Gln-Ala-Arg-7-aminomethylcoumarine (AMC) was rapidly cleaved by dasTMPRSS2, C-terminal to Arg, thereby releasing AMC product and enabling initial reaction velocities (*V_o_*) measurement within 60 seconds of enzyme addition (Fig. 3a). In Assay Buffer, dasTMPRSS2 had a *K_m_* of (200±80) μM, *V_max_* of (0.7±0.2) nmol min^−1^, *k_cat_* of (18±4)s^−1^, *k_cat_*/*K_m_* of (5.4±0.2) μM^−1^min^−1^ and specific activity at (0.22±0.03) μmol min^−1^mg^−1^ enzyme purified to apparent homogeneity (Fig. 3b). To our knowledge, this level of activity has not been achieved with any previously described recombinant TMPRSS2 enzyme^17–20^, and enzyme activity was unaffected by the presence of Ca^2+^, NaCl concentrations ranging 75-250 mM, EDTA, and tolerant of 2% (v/v) DMSO (Extended Data Fig. 4c) that is encouraging for use in high throughput inhibitor screening campaigns.

**Figure 3.**
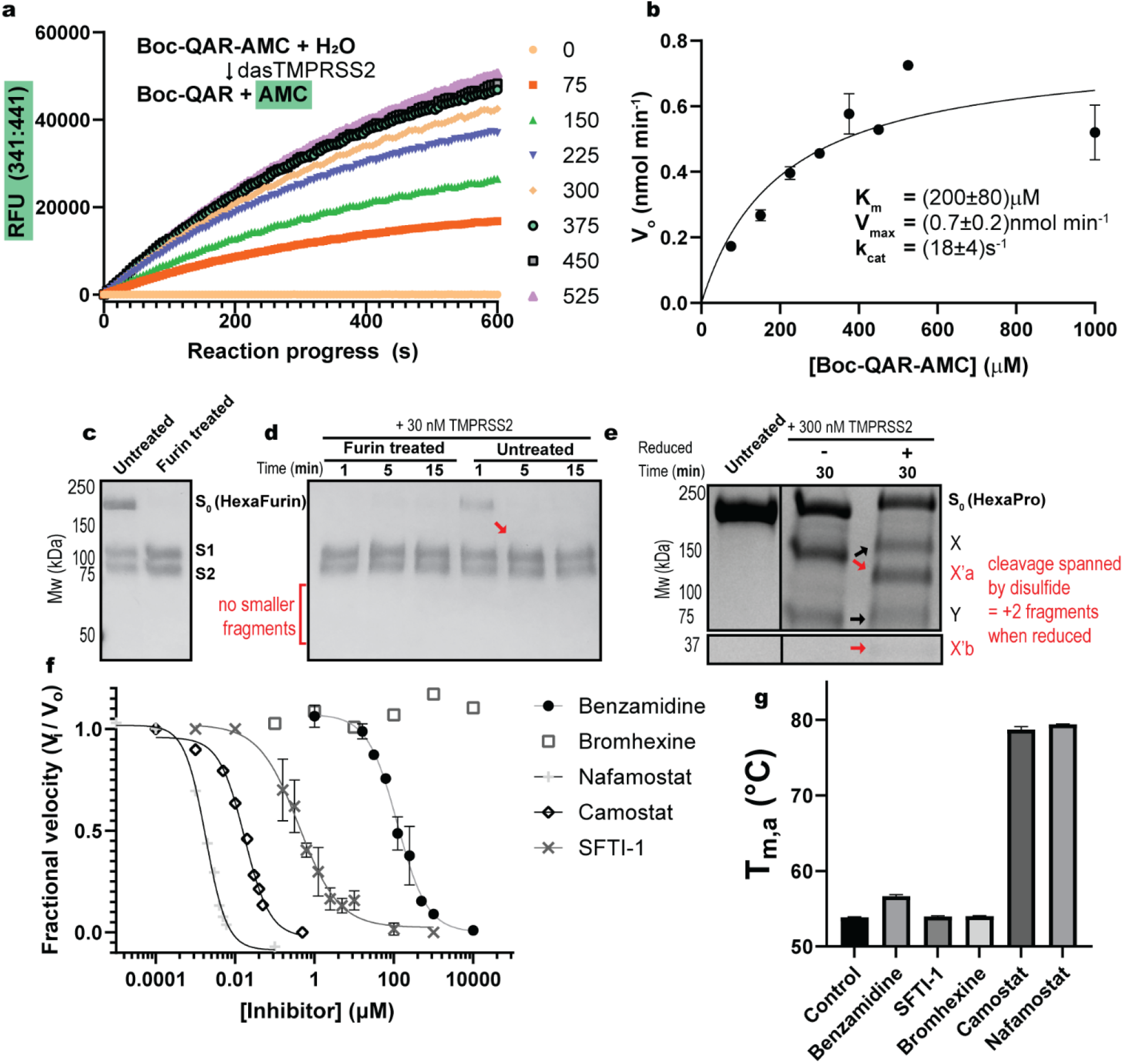
dasTMPRSS2 displays robust proteolytic activity towards peptide and SARS-CoV-2 S protein substrates. **a** The generic Boc-Gln-Ala-Arg-AMC fluorogenic peptide substrate is efficiently cleaved by dasTMPRSS2. Each progress curve was performed in quadruplet. **b** Michaelis-Menten plot of initial reaction velocities for kinetic parameter estimation after curve fitting in GraphPad. **c** S1/S2 intact S protein ectodomain (HexaFurin; S_0_) is partially cut at the S1/S2 site to produce S1 and S2 fragments and can be quantitatively converted by recombinant furin treatment over 16 hours. **d** The addition of 30 nM TMPRSS2 to furin treated HexaFurin produces no additional bands, but furin untreated HexaFurin is exhaustively cleaved at S1/S2 within 5 minutes. **e** dasTMPRSS2 cleaves HexaPro at two additional sites peripheral to S1/S2, with the first cleavage producing X and Y band fragments under non-reducing SDS-PAGE conditions. Reducing conditions reveal band fragments X’a and X’b derived from fragment X. **f** dasTMPRSS2 peptidase activity is blocked with varying potencies by clinical protease inhibitors, with no inhibition seen for bromhexine. All data are shown as mean ± s.e.m., *n* = 3 biological replicates **g** Apparent melting temperatures (as determined by differential scanning fluorimetry) are increased for benzamidine, camostat, and nafamostat at 1 μM concentration but are not increased by SFTI-1 or bromhexine. Samples were in triplicate.

### TMPRSS2 efficiently cleaves the SARS-CoV-2 S protein at the S1/S2 site in vitro

Cells expressing TMPRSS2 have been shown to efficiently cleave the SARS-CoV-1 S protein at S1/S2 (SLLR667↓) and multiple peripheral sites to induce the necessary conformational changes leading to virus-host fusion at the plasma membrane^21,22^ (Fig. 1a; Extended Data Fig. 5a). This extensive TMPRSS2 processing has also been linked to periplasmic shedding of the S1 fragment to act as an immune decoy in vivo^22^. For SARS-CoV-2, an acquired multibasic RRAR685↓ S1/S2 cleavage sequence was hypothesized to confer preferential cleavage by intracellular furin protease^23^, and was corroborated by studies showing that multibasic, peptidomimetic furin inhibitors prevented S1/S2 cleavage and attenuate infection in cellular models^24^. Further studies showed that these inhibitors are promiscuous and disable multiple surface-expressed proteases that process multibasic substrates in addition to furin, and more selective furin inhibitors cannot fully abrogate S activation^25^. Furthermore, furin-deficient cells can still generate S1/S2 cleaved virus, and propagation of SARS-CoV-2 in TMPRSS2-deficient cell lines results in a loss of the multibasic S1/S2 site^26^, attenuating viral infectivity towards TMPRSS2+ cells. We sought to characterize TMPRSS2’s proteolytic activity towards S1/S2 by incubating recombinant furin and/or dasTMPRSS2 with stabilized SARS-CoV-2 S protein ectodomain with S1/S2 knocked out (RRAR^685^->GSAS^685^; HexaPro construct) or with S1/S2 intact (denoted HexaFurin; Extended Data Fig. 5a; Fig. 3c). As expected from previous studies using recombinant, S1/S2 intact S protein, HexaFurin sustained partial S1/S2 cleavage during production in HEK293 cells due to endogenously expressed furin^27^ (Fig. 3c). Recombinant furin treatment converted the remaining intact HexaFurin to S1 and S2 band fragments with incubation over 16 hours, but was unable to cleave HexaPro (Fig. 3c; Extended Data Fig. 5b).

In contrast, using both the HexaFurin and HexaPro constructs, we observed that dasTMPRSS2 could cleave the S protein at 3 distinct sites with variable efficiency (Fig. 3d-e). HexaFurin was cleaved to only the S1 and S2 fragments within 5 minutes of dasTMPRSS2 addition (Fig. 3d), demonstrating the S1/S2 site was best recognized by TMPRSS2 across a minimal incubation. HexaPro, lacking S1/S2, was cleaved across 30 min to generate a larger 150 kDa band, denoted fragment X, and 80 kDa fragment Y when analyzed under non-reducing conditions (Fig. 3e). Reducing conditions revealed an additional cleavage site hidden within fragment X that is spanned by two cysteine residues participating in a disulfide bond, splitting fragment X into 120 kDa fragment X’a and 35 kDa fragment X’b. Exhaustive HexaPro treatment (120 min) completely converted fragment X into X’a and X’b (Extended Data Fig. 6d).

To visualize all of these cleavage sites simultaneously, we treated HexaFurin 30 min with dasTMPRSS2 and compared SDS-PAGE banding to a western blot using an antibody directed towards the S protein receptor binding domain (RBD; Extended Data Fig. 5c). At least 7 bands were observed on reducing SDS-PAGE and the western shows that both the S1 fragment as well as an S1-derived 50 kDa fragment contain the RBD. The banding patterns observed (S1/S2, X/Y, and X’a/X’b cleavages) are consistent with western blot studies monitoring SARS-CoV-1 S protein processing by TMPRSS2 that enables shedding of the S1 fragment^22^ to act as an immune decoy.

### Nafamostat rapidly acylates TMPRSS2 and slowly hydrolyzes

Nafamostat and camostat are serine protease inhibitors under investigation as anti-TMPRSS2 COVID-19 therapeutics (ClinicalTrial.gov identifiers NCT04583592, NCT04625114). Both are reactive esters that form the same slowly-reversible phenylguanidino covalent complex (evidenced in the enteropeptidase-camostat structure; PDB: 6ZOV) with the catalytic serine residue of trypsin-like serine proteases. Nafamostat and camostat dramatically increased the apparent melting temperature (*T_M,a_*) of dasTMPRSS2 by (25.5±0.1) and (24.8±0.3) °C, respectively, as measured by Differential Scanning Fluorimetry (DSF)^28^ (Fig. 3g) and was a key stabilizing feature to enable protein crystallization (Methods). Nafamostat demonstrated enhanced potency over camostat with IC_50_ values of (1.7±0.2) and (17±4) nM, respectively, with 5 min assay pre-incubation (Fig. 3f). However, IC_50_ values were time-dependent and required further kinetic interrogation to assess their divergent potencies (Extended Data Fig. 6b-c). Nafamostat was 40-fold more potent than camostat with respective *k*_inact_/*K*_i_ values of (0.024±0.006) and (0.00059±0.00003) s^−1^ nM^−1^. These results emphasize that single timepoint IC_50_ values are insufficient for evaluating mechanism-based, covalent inhibitors of this highly active protease in SAR studies. As previously identified for matriptase, the nafamostat leaving group, 6-amidino-2-napthol, fluoresces and can be used as a sensitive burst titrant to calculate the concentration of active protease (Extended Data Fig. 4e-f; Methods)^19^. The half-life of the phenylguanidino acyl-enzyme complex was (14.7±0.4) hours as measured by the gradual rescue of dasTMPRSS2 peptidase activity after stoichiometric acylation with nafamostat (Extended Data Fig. 3e-f). Impressively, stoichiometric amounts of nafamostat completely blocked dasTMPRSS2-mediated HexaPro activation over 2 hours (Extended Data Fig. 6d) and are consistent with this drug’s ability to potently block SARS-CoV-2 pseudovirus entry to TMPRSS2+ Calu-3^29^ and Caco-2^5,20^ lung cells.

Non-covalent trypsin-like serine protease inhibitors benzamidine and sunflower trypsin inhibitor-1 (SFTI-I) were less potent with respective IC_50_ values of (120±20) μM and (0.4±0.2) μM (Fig. 3f), and *K_i_* values of (80±10) μM and (0.4±0.2) μM (Extended Data Fig. 7a-b). 6-amidino-2-napthol also disabled dasTMPRSS2 activity with an IC_50_ of (1.6±0.5) μM and *K_i_* of (1.1±0.3) μM (Extended Data Fig. 7a). Bromhexine hydrochloride, another agent under investigation for anti-TMPRSS2 COVID-19 therapy^30,31^, showed no inhibition in either the peptidase or HexaPro cleavage assay formats (Extended Data Fig. 7c-d), corroborating reports of its ineffectiveness in blocking SARS-CoV-2 pseudovirus entry^32^ and further underscores the need for novel, selective TMPRSS2 inhibitors.

### Future prospects

We have produced and characterized a source of TMPRSS2 enzyme that will enable rapid inhibitor development as antivirals and thorough molecular interrogation of coronavirus and influenza virus activation. Although nafamostat potently neutralizes TMPRSS2 activity, it is non-selective and disables trypsin-like serine proteases involved in coagulation such as plasmin, FXa, and FXIIa, as well as other TTSPs through its generic arginine-like engagement with the S1 subsite^19,33,34^. Furthermore, nafamostat requires continuous intravenous infusion to approach therapeutic concentrations for COVID-19 owing to its short biological half-life of 8 minutes (NCT04418128; NCT04473053). These features, although undesirable as a selective therapeutic, make nafamostat an extremely useful and sensitive reagent for *in vitro* kinetic characterization of trypsin-like proteases, and sufficiently stabilized our protease for crystallization and structural determination. Nevertheless, selective and biologically stable drugs for TMPRSS2 must be explored, and may be achieved through inhibitors engaging the more TMPRSS2-specific S2, S3, and S4 subsites identified in our crystal structure.

We observed no electron density for the LDLR-A domain of TMPRSS2, despite a similar construct design to that which afforded the TMPRSS13 crystal structure (PDB: 6KD5). The LDLR-A domain of TTSPs is responsible for tethering the protease to the plasma membrane and most TTSPs have a conserved ability to bind calcium. Interestingly, a key Asp residue in TMPRSS13 involved in calcium chelation is absent in human and other mammalian TMPRSS2 proteins, substituted instead with His or Gln residues (Extended Data Fig. 8). These data suggest that TMPRSS2 may have lost the ability to bind calcium at this site.

Our demonstration that TMPRSS2 can cleave the multibasic S1/S2 site of the S protein suggests that instead of conferring furin dependence, the virulent properties of this site may derive from promiscuous recognition and cleavage by airway-expressed TTSPs, which is supported by the demonstrated roles that TMPRSS4^35,36^, TMPRSS11d^20,37,38^, and TMPRSS13^20,37^, which colocalize with ACE2^36^, play in enabling SARS-CoV-2 infection across various tissues.

Our characterization of dasTMPRSS2 did not reveal an obvious mechanism by which the native, membrane-bound enzyme could be autoproteolytically processed peripheral to the activation motif and thereby shed as a soluble enzyme into the extracellular space. However, studies using TMPRSS2-specific antibodies have reported detection of a secreted enzyme product in prostate sera that is expected to play a functional role in pericellular activation^39^. Due to the disulfide-linked nature of activated TMPRSS2, former studies may have mischaracterized the catalytic subunit as a shed SP domain when it would instead resolve to the intact species under non-reducing conditions (Fig. 1b). Thus, a biochemical characterization of these secreted species is required to interpret their activation status and subunit organization, as an active, shed form of TMPRSS2 in the extracellular milieu would have profound pathobiological and therapeutic targeting implications.

## ACKNOWLEDGEMENTS

We thank Jason McLellan for generously providing the SARS-CoV-2 S protein construct plasmids, Irene Chau for assistance with DSF, and Shih-Ting Tseng for preparation of Figure 1 graphic art. This work was supported by BC Leadership Chair in Functional Cancer Imaging to FB, the Canada Excellence Research Chair to SS, and a Mitacs Accelerate Internship to BF. This work is based upon research conducted at the Northeastern Collaborative Access Team beamlines, which are funded by the National Institute of General Medical Sciences from the National Institutes of Health (P30 GM124165). The Eiger 16M detector on 24-ID-E beam line is funded by a NIH-ORIP HEI grant (S10OD021527). This research used resources of the Advanced Photon Source, a U.S. Department of Energy (DOE) Office of Science User Facility operated for the DOE Office of Science by Argonne National Laboratory under Contract No. DE-AC02-06CH11357. The Structural Genomics Consortium is a registered charity (no: 1097737) that receives funds from AbbVie, Bayer AG, Boehringer Ingelheim, Genentech, Genome Canada through Ontario Genomics Institute [OGI-196], the EU and EFPIA through the Innovative Medicines Initiative 2 Joint Undertaking [EUbOPEN grant 875510], Janssen, Merck KGaA (aka EMD in Canada and US), Pfizer, Takeda and the Wellcome Trust [106169/ZZ14/Z].

## Author Information

These authors contributed equally: Bryan J. Fraser, Serap Beldar

## Author Contributions

F.B., C.H.A., L.H., and S.S. provided project supervision; B.J.F., S.B., A.S., C.H.A., and F.B. conceived the project; B.J.F., S.B., A.S., C.H.A., and L.H. designed the experiments; Y.L. cloned TMPRSS2 protein constructs for expression and Y.L. and D.M cloned SARS-CoV-2 S protein constructs for expression; A.S. and A.H. produced TMPRSS2 protein in insect cells; A.S., A.H., D.M., K.L. produced SARS-CoV-2 S protein in HEK cells; B.J.F., S.B., A.S., A.H., D.M., and K.L. purified recombinant proteins; B.J.F., S.B., and L.H. crystallized TMPRSS2 and L.H., S.B., and B.J.F. collected diffraction data; L.H. solved the crystal structure; B.J.F., S.B., and L.H. performed bioinformatic and structural analyses; B.J.F., D.K., R.W., and R.T. performed fluorogenic peptidase activity and inhibitor potency assays and B.J.F. analyzed kinetics; D.K. synthesized, purified, and characterized SFTI-1 peptide; B.J.F., D.K., and S.B. managed inhibitor compound libraries; B.J.F., D.M., and R.T. performed gel-based S protein digestion assays; S.B. and B.J.F. performed DSF assays; B.J.F., S.B., L.H., R.T., and D.M. prepared figures; B.J.F., S.B., L.H., C.H.A., F.B., A.S., A.H., and Y.L. wrote the manuscript.

## Competing interests

The authors declare no competing interests.

## Data Availability

The coordinates and structure of the phenylguanidino TMPRSS2 acyl-enzyme complex have been deposited in the PDB with accession number 7MEQ on April 7, 2021, and released on April 21, 2021. Any other relevant data are available from the corresponding authors upon reasonable request.

## METHODS

### Construct design and cloning

A construct encoding residues 109-492 comprising soluble TMPRSS2 ectodomain was amplified by two PCR fragments (Addgene plasmid# 53887) and subcloned into the pFHMSP-LIC C donor plasmid by LIC method. The final construct contained a N-terminal honeybee melittin signal sequence peptide and C-terminal His_8_-tag (Fig.1b). Mutations targeting the activation sequence SSQSR255↓IVGGE (arrow indicates the cleavage site) were implemented to replace the SRQSR255 residues with an enteropeptidase-cleavable DDDDK255 graft with two sets of primer pairs (Extended Data Table 1) generating mutations for S251D/R252D/Q253D/S254D/R255K. Plasmid transfer vector containing the TMPRSS2 gene was transformed into *Escherichia coli* DH10Bac cells (Thermo Fisher; Cat# 10361012) to generate recombinant viral bacmid DNA. Sf9 cells were transfected with Bacmid DNA using JetPrime transfection reagents (PolyPlus Transfection Inc.; Cat# 114-01) according to the manufacturer’s instructions, and recombinant baculovirus particles were obtained and amplified from P1 to P2 viral stocks. Recombinant P2 viruses were used to generate suspension culture of baculovirus infected insect cells (SCBIIC) for scaled-up production of TMPRSS2.

The SARS-CoV-2 Spike ectodomain HexaPro construct was a gift from J. McLellan^1^, and the S1/S2 site was restored (GSAS^685^->RRAR) through site-directed mutagenesis with primers in Extended Data Table 1 (HexaFurin construct).

### Baculovirus mediated dasTMPRSS2 protein production in Sf9 insect cells

Sf9 cells were grown in I-Max Insect Medium (Wisent Biocenter; Cat# 301-045-LL) to a density of 4×10^6^ cells/mL and infected with 20 mL/L of suspension culture of baculovirus infected insect cells prior to incubation on an orbital shaker (145 rpm, 26 °C).

### dasTMPRSS2 protein purification

Cell culture medium containing the final secreted protein product AA-[TMPRSS2(109-492)]-EFVEHHHHHHH was collected by centrifugation (20 min, 10 °C, 6,000 x g) 4-5 days post-infection when cell viability dropped to 55 - 60%. Media was adjusted to pH 7.4 by addition of concentrated PBS stock, then supplemented with 15 mL/L settled Ni-NTA resin (Qiagen) at a scale of 12 L. Three batch Ni-NTA purifications were used to capture protein, with each round requiring shaking in 2L flasks for 2 hours at 16 °C (110 rpm), harvesting by centrifugation (5 min 1,000 x g), then transferred to a gravity flow column. Beads were washed with 3 column volumes (CVs) ice-cold PBS prior to elution with PBS supplemented with 500 mM imidazole. Elution samples were concentrated to 4.5 mg/mL using 30 kDa MWCO Amicon filters and overnight zymogen activation was achieved by dialyzing protein 1:1000 against Assay Buffer (25 mM Tris pH 8.0, 75 mM NaCl, 2 mM CaCl_2_) at room temperature in the presence of recombinant enteropeptidase (NEB) at 13 U enzyme per mg TMPRSS2 zymogen. The following day, activated samples were exchanged to SEC buffer (50 mM Tris pH 7.5, 250 mM NaCl), spun down at 17,000 x g, then loaded to a Superdex 75 gel filtration column. Fractions spanning the dominant peak eluting at 80 mL (Extended Data Fig. 1) were evaluated for appropriate banding on reducing SDS-PAGE prior to pooling and concentrating. For dasTMPRSS2 enzyme samples, 2 μL aliquots of 10,000x enzyme assay stocks (32 μM) were prepared by concentrating protein to 1.6 mg/mL in Enzyme Buffer (50 mM Tris pH 7.5, 250 mM NaCl, 25% glycerol), then flash-frozen in liquid nitrogen and stored at –80 °C until thawed immediately prior to use for each enzyme assay in order to minimize autoproteolysis and maintain reproducible enzyme concentrations.

### HexaPro and HexaFurin production and purification

Expi293F cells (Life Technologies Cat. # A1435102) were transiently transfected with expression plasmid encoding HexaPro/Furin using FectoPro transfection reagent (Polyplus-transfection® SA, Cat. #116-010) with 5 mM sodium butyrate being added at the time of transfection (Sigma, Cat. # 303410). After 4-5 days post-transfection time cells culture was harvested, supernatant cleared by centrifugation, and the pH was adjusted by adding 10x Buffer (50 mM Tris pH 8.0, 150 mM NaCl). Secreted protein was captured by two round of batch absorption (BA) with 4 mL/L of pre-equilibrated Ni Sepharose beads (GE Healthcare, Cat #17-5318-01). The bound beads were transferred to gravity flow column and sequentially washed with 30 CVs Wash Buffer (50 mM HEPES 7.5, 300 mM NaCl, 5% glycerol), followed by Wash Buffer supplemented with 25 mM imidazole. Protein was eluted in Elution Buffer (Wash buffer with 250 mM imidazole) and concentrated using Amicon™ Ultra Centrifugal Filter Units, 15 mL, 100 kDa (Millipore Sigma™ Cat# UFC910024) prior to size-exclusion chromatography purification using a Superose 6 Increase 10/300 GL (GE Healthcare Cat # 29-0915-96), in a buffer composed of 20 mM HEPES pH 7.5, 200 mM NaCl.

### Protein crystallization and structural determination

After size-exclusion purification of activated dasTMPRSS2, samples were pooled and concentrated to 2 mg/mL. Protein was treated with 3:1 nafamostat:dasTMPRSS2 for 10 minutes at room temperature and exchanged into Assay Buffer supplemented with 3:1 nafamostat using 4 spin cycles in 30 kDa Amicon MWCO filters (14,000 rpm, 15 min, 4 °C) to remove low Mw autolytic fragments from the 42 kDa enzyme (Extended Data Fig. 1b). Acylated enzyme was then concentrated to 8 mg/mL and centrifuged (14,000 rpm, 10 min, 4 °C) prior to automated screening at 18 °C in 96-well Intelliplates (Art Robin) using the Phoenix protein crystallization dispenser (Art Robbins). Protein was dispensed as 0.3 μL sitting drops and mixed 1:1 with precipitant. The RedWing and SGC precipitant screens were tested and amorphous, non-diffracting crystals were consistently produced when grown over 30% Jeffamine ED-2001 (Hampton Research) with 100 mM HEPES pH 7.0. To acquire a diffraction quality crystal, acylated dasTMPRSS2 was treated with 50 U PNGase F (NEB; 37 °C for 45 min) to trim N-glycan branches, then centrifuged (14,000 rpm, 4 °C, 10 min) prior to setting 2 μL hanging drops with 1:1 protein: precipitant and grown for 10 days. Crystals were then cryo-protected using reservoir solution supplemented with ~5% (v/v) ethylene glycol, and cryo-cooled in liquid nitrogen. X-ray diffraction data were collected on the beamline 24-ID-E at the Advanced Photon Source (APS). Data were processed with XDS^2^. Initial phases were obtained by molecular replacement in Phaser MR^3^, using (PDB: 1Z8G) as a starting model. Model building was performed in COOT^4^ and refined with Buster^5^. Structure validation was performed in Molprobity^6^. Data collection and refinement statistics are summarized in Extended Data Table 2.

### Gel electrophoresis and western blotting

SDS-PAGE was carried out with 15 μL Mini-Protean (BioRad) or 60 μL Novex Wedgewell (Invitrogen) 4-20% Tris-Glycine gels for 30 min under constant voltage at 200V. Protein samples were mixed with 4x Laemelli buffer (BioRad) and subjected to differential reducing (± 5 mM β-mercaptoethanol; Gibco), then boiling at 95 °C for 5 min in order to probe the covalent nature of protein complexes and subunits. The Precision Plus Protein marker (BioRad) was used as a standard.

For SARS-CoV-2 RBD western blotting, SDS-PAGE was carried out as described, followed by wet transfer in Transfer Buffer (25 mM Tris pH 8.3, 192 mM glycine, 20% MeOH (v/v)) to PVDF membrane (80 V, 53 min, 4 °C). Membranes were incubated in Blocking Buffer (5% skim milk in TBST) for 1 hr at room temperature, washed 5x with TBST, then probed overnight with 1/3000 mouse anti-RBD primary mAb (Abcam ab277628) solution at 4 °C. Membranes were then washed 5x with TBST and probed with 1/5000 FITC-labelled goat anti-mouse IgG secondary pAb (Abcam ab6785) and imaged for fluorescence on the Typhoon FLA7000 biomolecular imager (GE healthcare).

### Enzyme peptidase and inhibition assays

Peptidase assays with fluorogenic Boc-Gln-Ala-Arg-AMC substrate (Bachem Cat # 4017019.0025) were performed in 96-well plates (Greiner Fluotrak) at 200 μL reaction volumes in a FlexStation microplate reader (Molecular Devices) at 24 °C. Fluorescence was monitored with the fastest kinetic read settings across 5 minutes at 341:441 nm excitation:emission and converted to a product AMC concentration using standard curves at each substrate concentration to correct for the inner-filter effect^7^ (Extended Data Fig. 4d). All assays contained 2% (v/v) DMSO and initial reaction velocities were tabulated over the linear portion of the first 60 seconds of progress curves.

To determine Michaelis Menten kinetic parameters, 50 μL 4x enzyme stock was added through automated addition to microplates containing 150 μL substrate (0.5-1000 μM) in triplicate and initial reaction velocities were plotted against substrate concentration and curve fit using GraphPad Prism.

Half-maximal inhibitor (IC_50_) potencies of nafamostat mesylate (MedChemExpress Cat # HY-B0190A), camostat mesylate (MedChemExpress Cat # HY-13512), benzamidine HCl (Sigma Cat # 434760-25G), bromhexine HCl (SelleckChem Cat # S2060), and SFTI-1 were initially determined by pre-incubating dasTMPRSS2 with inhibitor at concentrations ranging from 0.1 nM – 100 μM for 10 min, then enzyme-inhibitor mixes were added to substrate through automated addition. Then, 7 inhibitor concentrations spanning three orders of magnitude across the IC_50_ value were used and inhibitor reaction velocities were normalized to uninhibited enzyme and plotted as one-site dose response curves in GraphPad. The apparent Ki (Ki*) of classical competitive trypsin-like serine protease inhibitors benzamidine and SFTI-1 were determined using Equation 1,

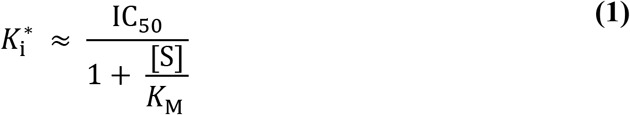

Where [S] is the concentration of substrate Boc-QAR-AMC and *K*_M_ is the Michaelis constant.

### Time-dependent IC_50_ measurement and *k*_inact_/*K*_i_ determination

Camostat IC_50_ curves were generated using 7 concentrations of inhibitor ranging 0.1-1000 nM inhibitor and nafamostat between 0.01-100 nM with a DMSO control as described. The time dependence of inhibitor potencies was measured by using Flexstation Flex kinetic reads that automatically transferred dasTMPRSS2-inhibitor mixes to substrate wells at the indicated pre-incubation timepoints (Extended Data Fig. 6c). For the 10s timepoint, a kinetic read was performed after manual addition of enzyme, followed by substrate, using a multichannel pipette. Kinetic parameters Kiapp and kinact were determined with the simplified Equations 2 and 3, respectively, assuming a one-step kinetic inhibition mechanism, A^8^.

#### Mechanism A

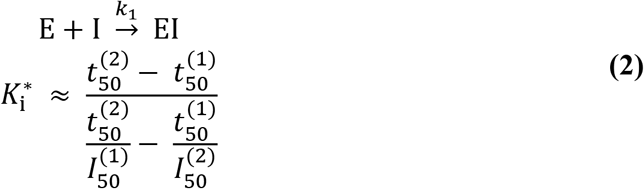

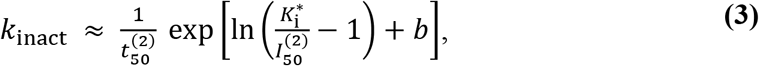

where the conversion factor *b* = 0.558 is applied for concentrations in μM and time in s.

### Active site quantification of dasTMPRSS2

The acylation of dasTMPRSS2 by nafamostat and concomitant production of the fluorogenic 6-amidino-2-napthol leaving group was measured by incubating serial log2 enzyme dilutions from 6.4-0.8 nM with excess (10 μM) nafamostat, similar to previous efforts with matriptase^9^ (Extended Data Fig. 4e-f). Microplate reading at 320: 490 nm excitation:emission were used to calculate the number of dasTMPRSS2 active site residues and calibrate peptidase activity and inhibition assays.

### Nafamostat inhibition half-life

The half-life of the phenylguanidino acyl-enzyme complex after nafamostat treatment was measured for dasTMPRSS2 using methods established for camostat with enteropeptidase^10^. Briefly, dasTMPRSS2 (3.2 μM) was mixed with slight excess nafamostat (5 μM or DMSO control) and incubated at room temperature for 20 minutes. After incubation, unbound nafamostat was removed by passage and 3x washes in a 3 kDa MWCO Amicon filter centrifuged at maximum speed. Acylated and untreated dasTMPRSS2 samples were then transferred in quadruplet to a microplate containing either 125 μM or 250 μM substrate (final concentration of 3.2 nM enzyme). Fluorescent reads were carried out immediately, analogous to IC_50_ assays, but across a period of 8 hours. The acylated traces were fit to a one-phase association exponential in GraphPad to derive the half-life for activity recovery, normalized to the uninhibited initial reaction velocity.

### Differential Scanning Fluorimetry

Apparent melting temperature (*T_M,a_*) shifts were measured for various dasTMPRSS2-inhibitor coincubations using SYPRO Orange dye (Life Technologies; cat. S-6650) and monitoring fluorescence at 470:510 nm excitation: emission using the Light Cycler 480 II (Roche Applied Science). Samples were prepared in triplicate in 384 well plates (Axygen; Cat# PCR-384-C; Cat# UC500) at a final volume of 20 μL containing 0.05 mg/mL dasTMPRSS2, 1 μM compound or vehicle control, and 5X SYPRO Orange. Thermal melt curves were generated between 25 °C to 95 °C at a gradient of 1 °C /min and plots prepared with the DSFworld application^11^ for *T_M,a_* determination.

### SARS-CoV-2 S protein activation and inhibition

Recombinant SARS-CoV-2 S protein constructs HexaFurin and HexaPro were concentrated to 0.5 mg/mL in Assay Buffer and incubated with the indicated concentrations of furin protease (NEB) or dasTMPRSS2. Digestions took place over 16 hours for furin and from 5-120 minutes for dasTMPRSS2. Furin digestions were terminated by the addition of 4 mM EDTA whereas dasTMPRSS2 digestions were terminated with 5 μM nafamostat, then SDS-PAGE samples were immediately prepared with the addition of 4X SDS-PAGE loading buffer and boiled for 5 min at 95 °C. 4 μg S protein were loaded per well under each conditions and gels visualized by Coomassie blue staining. For anti-RBD western blotting, 2 μg S protein were loaded per well. For cleavage inhibition assays, dasTMPRSS2 diluted to 320 nM in Assay Buffer was pre-incubated 15 minutes with inhibitor (final 1% DMSO (v/v)) or DMSO control, then assays were started by transfer of enzyme:inhibitor mixes to S protein. S protein:protease mixtures were incubated at room temperature for 2 hours with nafamostat and 30 minutes for bromhexine.

### Multiple Sequence Alignments

Multiple sequence alignments were prepared to compare the human TTSP family members and TMPRSS2 mammalian orthologs. Human TTSP FASTA sequences (isoform 1) were accessed from UniProt and TMPRSS2 orthologs identified with UniProt BLAST. Sequences were aligned with Clustal Omega^12^ and annotated with ESPript 3.0^13^.

### Protein Visualization and Property Calculation

The structure of dasTMPRSS2 was inspected and compared to other TTSPs using PyMol (Schrodinger) and the Molecular Operating Environment (MOE; Chemical Computing Group) software suite. The exposed hydrophobic patches of TMPRSS2 were calculated using the MOE Protein Patch Analyzer tool^14,15^.

## Notes

### Competing Interest Statement

The authors have declared no competing interest.

