## Extended Data for "Structure, activity and inhibition of human TMPRSS2, a protease implicated in SARS-CoV-2 activation"

**Extended Data Table 1. Mutagenesis primer sequences**

| LIC primer purpose | | Primer sequence (5′ to 3′) |
|---|---|---|
| TMPRSS2 SRQSR255↓IVGGE 1st PCR | Fwd | tacatctatgcggccgctatgggcagcaagtgctccaac |
| | Rev | cttgtcgtcgtcgtcgttcaagttgaccccgcaggc |
| TMPRSS2 SRQSR255↓IVGGE 2nd PCR | Fwd | gacgacgacgacaagatcgtgggcggcgagagcgcgctc |
| | Rev | ggtgctcgacgaattcgccgtctgccctcatttgtc |
| HexaPro GSAS685->RRAR 1st PCR | Fwd | tctcaggccgagttcggtaccgccaccatgttcgtgttcctggtgctcctgcctctgg |
| | Rev | agaacgagctcgtctagggctgtttgtctgggtctggtag |
| HexaPro GSAS685->RRAR 2nd PCR | Fwd | cctagacgagctcgttctgtagctagccagagcatcatcgcctac |
| | Rev | tcggggatgtatccggatccctgctcgtatttgccgagctcctgcaggtcg |

**Extended Data Table 2. Data collection and refinement statistics**

| | dasTMPRSS2 pre-treated with Nafamostat (PDB ID: 7MEQ) |
|---|---|
| **Data collection** | |
| Space group | P 1 $2_1$ 1 |
| Cell dimensions | |
| *a*, *b*, *c* (Å) | 59.45, 51.42, 64.35 |
| α, β, γ (°) | 90.00, 91.53, 90.00 |
| Resolution (Å) (highest resolution shell) | 43.08 - 1.95 (2.00 -1.95) |
| $R_{merge}$ | 0.067 (0.676) |
| $I / \sigma I$ | 8.9 (1.5) |
| Completeness (%) | 97.6 (98.9) |
| Redundancy | 4.4 (4.3) |
| **Refinement** | |
| Resolution (Å) | 40.17 - 1.95 |
| No. reflections | 27797 |
| $R_{work}$ / $R_{free}$ | 19.2/22.5 |
| No. atoms | 2640 |
| Protein | 2482 |
| Ligand | 12 |
| Waters | 118 |
| *B*-factors | 54.1 |
| Protein | 54.1 |
| Ligand | 49.5 |
| Waters | 46 |
| R.m.s. deviations | |
| Bond lengths (Å) | 0.01 |
| Bond angles (°) | 1.04 |

*Values in parentheses are for highest-resolution shell.

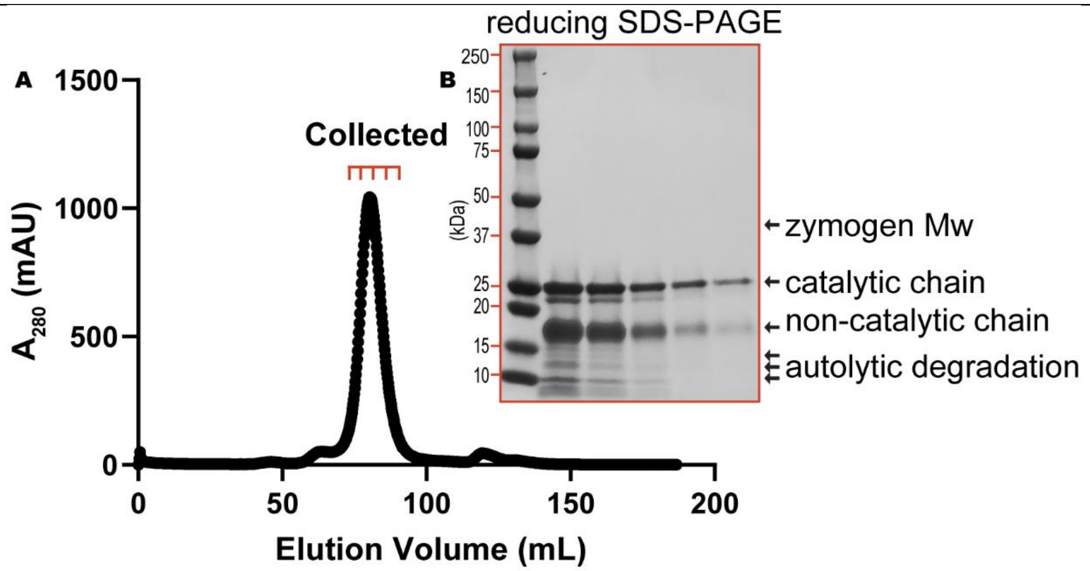

**Extended Data Fig. 1 Size-exclusion chromatography purification of crude, activated dasTMPRSS2. a** Size-exclusion chromatography profile of crude dasTMPRSS2 after overnight activation with recombinant enteropeptidase, with fractions collected around the dominant 80 mL peak. **b** Reducing SDS-PAGE (5% β-mercaptoethanol) analysis of collected fractions indicating complete zymogen activation and minor autolysis of concentrated fractions.

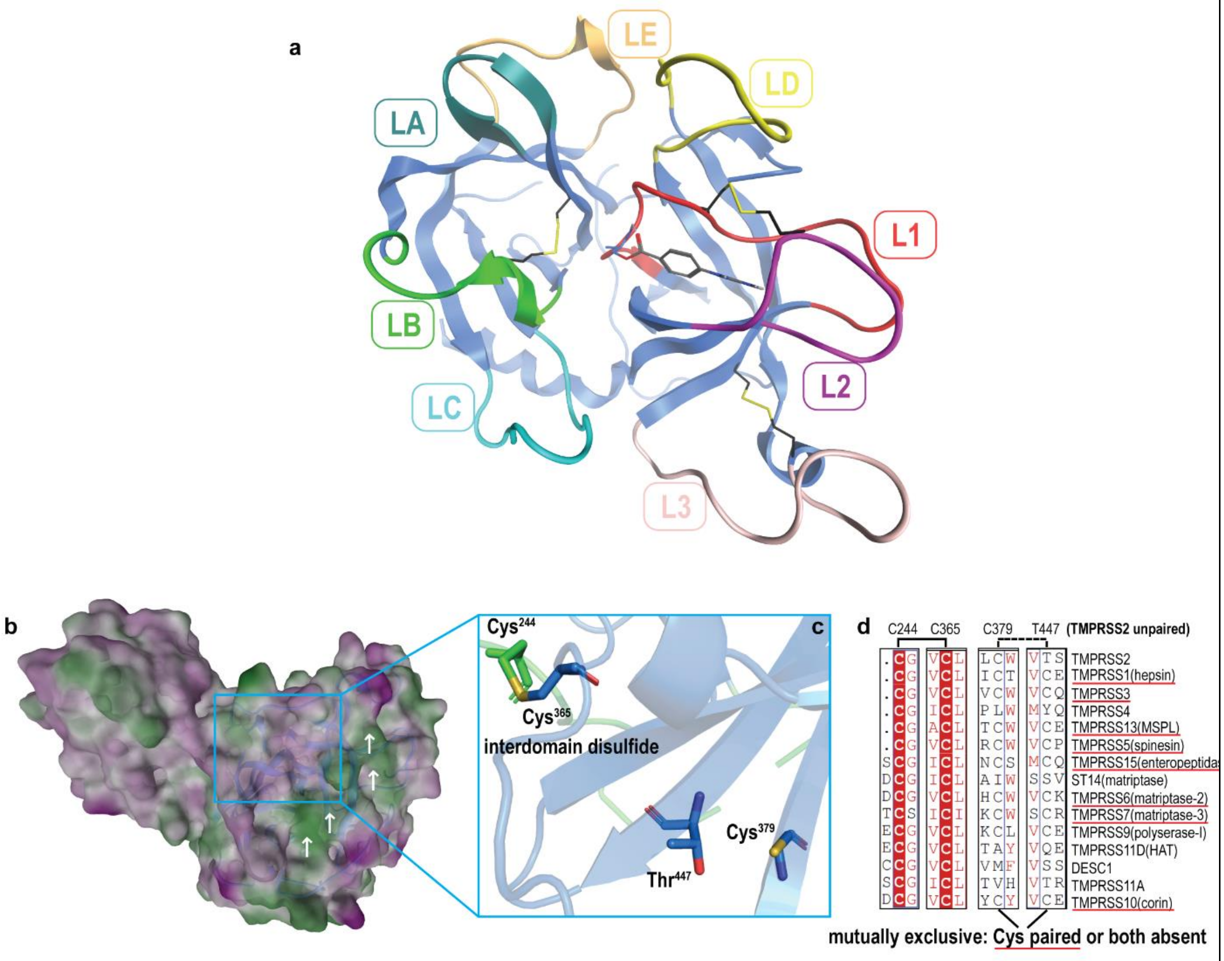

**Extended Data Fig. 2 Molecular recognition of TMPRSS2 substrates is mediated through surface loops and potentially through an unpaired cysteine residue. a** The substrate binding face of trypsin-like proteases makes use of 3 disulfides (yellow sticks) and 8 loops, Loops 1-3 (L1-L3) and LA-LE, to confer protein substrate specificity. Nafamostat (grey sticks) covalently bound to the catalytic Ser441 engages the S1 subsite of TMPRSS2 with residues from L1 and L2. **b** MOE protein patch analysis of TMPRSS2 ectodomain reveals a 360 Å$^2$ hydrophobic patch (green) highlighted with white arrows. **c** The unpaired Cys379 residue is adjacent to the hydrophobic patch and in close proximity to the highly conserved Cys244-Cys365 interdomain disulfide. **d** A multiple sequence alignment of the TTSP family identifies TMPRSS2 as uniquely possessing an unpaired cysteine residue in the SP domain.

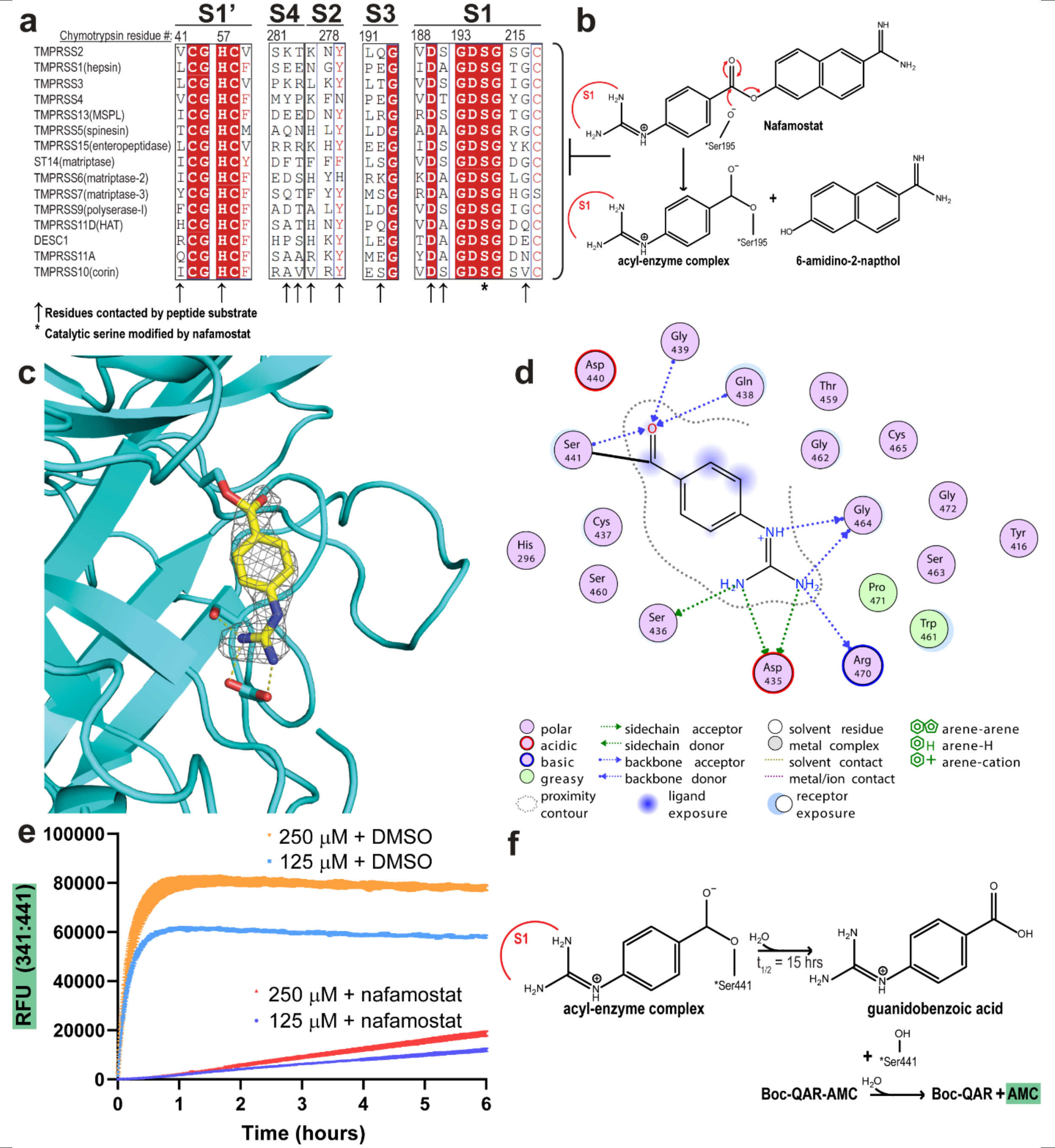

**Extended Data Fig. 3 Nafamostat acylation and deacylation of dasTMPRSS2**. **a** Multiple sequence alignment of the TTSP family, demonstrating high conservation of the S1 protease subsite that is utilized by nafamostat to acylate the catalytic serine residue, Ser195 (denoted with an asterisk). The S2-S4 protease subsites show greater variability than S1 and confer divergent substrate specificity within the TTSP family. **b** Nafamostat acts as a suicide substrate, forming a phenylguanidino acyl-enzyme complex with the catalytic serine residue, and produces a 6-amidino-2-napthol side product. **c** The mFo-DFc electron density omit-map of phenylguanidino in nafamostat-treated dasTMPRSS2 displayed as a grey mesh and contoured at 2.5σ.**d** Detailed nafamostat-dasTMPRSS2 interactions within the S1 protease subsite. **e** Recovery of dasTMPRSS2 peptidase activity (datapoints shown as mean ± s.e.m., *n* = 4 biological replicates) after acylation with nafamostat via (**f**) hydrolysis of the phenylguanidino acyl-enzyme complex to alleviate the catalytic Ser441.

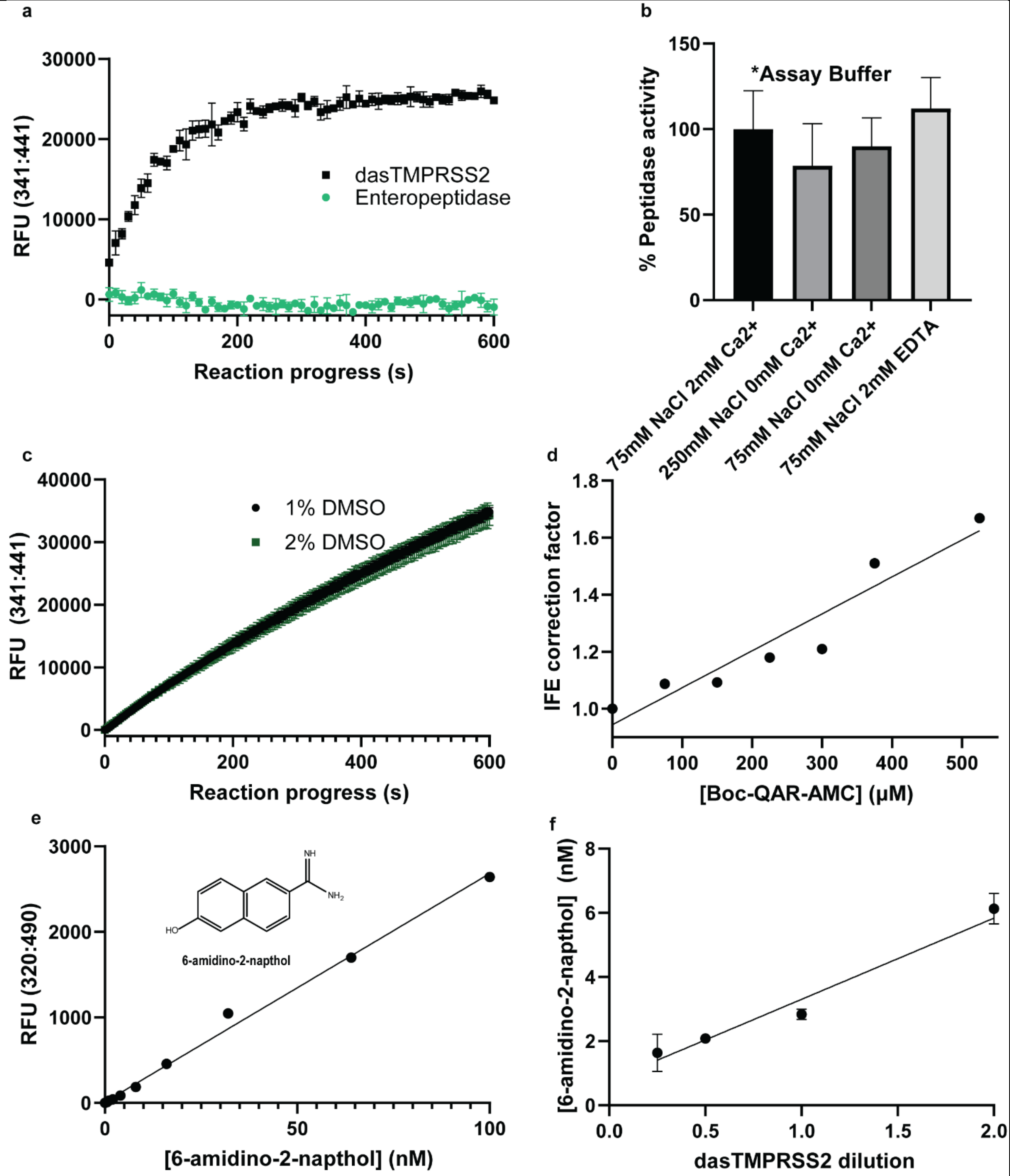

**Extended Data Fig. 4 Development of a robust and sensitive TMPRSS2 peptidase activity assay.** **a** Reaction progress curve for 5 µM Boc-QAR-AMC substrate and 3 nM dasTMPRSS2 or 0.03 U enteropeptidase (NEB). No proteolytic activity was observed for enteropeptidase, confirming that protetolytic activity is TMPRSS2-mediated. Data are shown as mean ± s.e.m., *n* = 3 biological replicates **b** Modified peptidase assay conditions in 25 mM Tris pH 8.0 show no significant differences when [NaCl] or [$Ca^{2+}$] is modified. Reaction velocities are relative to Assay Buffer (starred). **c**

Reaction progress curve for 100 µM Boc-QAR-AMC with final assay conditions containing 1% or 2% DMSO, demonstrating no loss in relative activity with increased DMSO. Data are shown as mean ± s.e.m., $n$ = 4 technical replicates **d** Required IFE correction factors for Michaelis Menten plot development (Fig. 2a-b) fit to a least-squares linear regression. **e** Fluorescence standard curve of the nafamostat leaving group, 6-amidino-2-napthol. **f** dasTMPRSS2 enzyme quantification after incubation with 1 µM nafamostat and concomitant production of 6-amidino-2-napthol. The indicated dilutions provided an estimated assay concentration of 3.2 nM enzyme for use in enzyme inhibition screens.

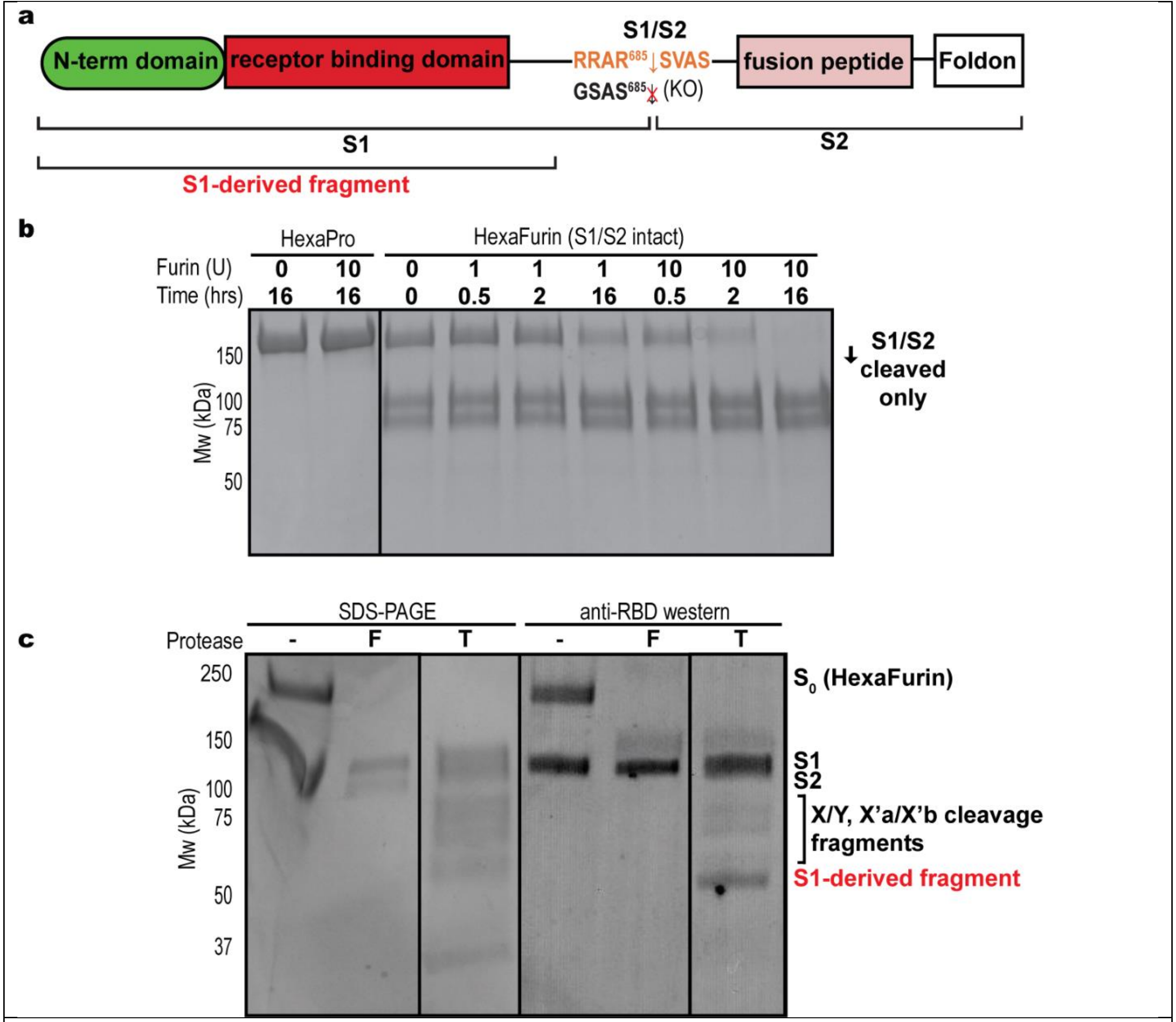

**Extended Data Fig. 5 Proteolytic processing of SARS-CoV-2 S protein ectodomain. a** The S protein is cut at the S1/S2 site by host proteases to release subunit 1 (S1) containing the N-terminal and receptor binding domains (RBD), and S2 containing virus-host membrane fusion machinery. An additional TMPRSS2 cleavage site within S1 appears to release a fragment containing RBD. **b** Recombinant furin (NEB enzyme units in parentheses) exclusively cuts the S protein at S1/S2 over a period of 16 hours at room temperature. **c** Exhaustive furin (F) treatment exclusively produces the S1 band fragment when analysed with an anti-RBD western blot whereas TMPRSS2 (T) treatment over 30 minutes produces S1, an S1-derived fragment containing the RBD, as well as numerous SDS-PAGE bands that are a combination of S1/S2, X/Y, and X'a/X'b cleavage events. SDS-PAGE and western blotting results were consistent with *n* = 3 biological replicates.

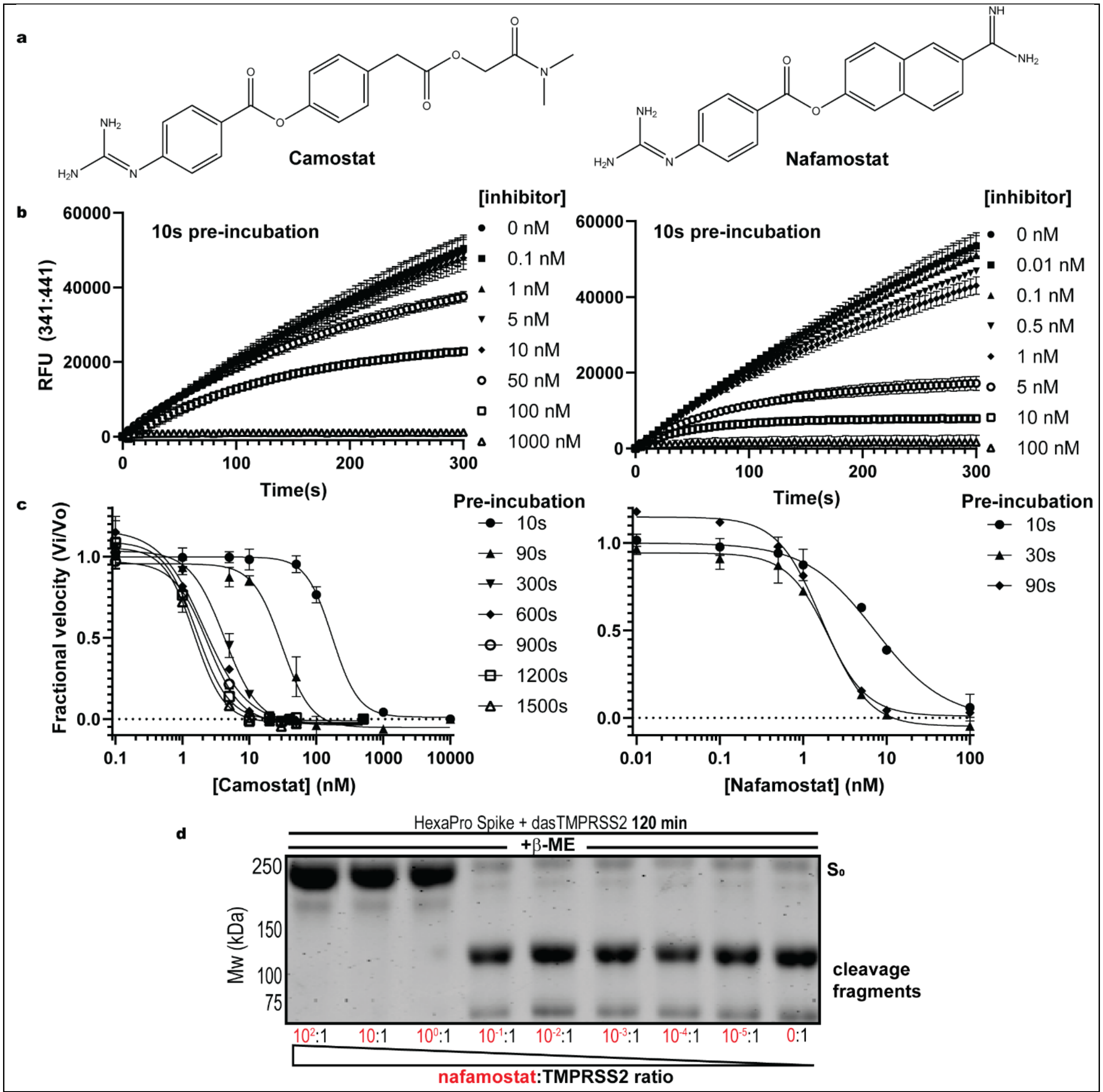

**Extended Data Fig. 6 Time-dependent dasTMPRSS2 inhibition by camostat and nafamostat. a** chemical structures of camostat and nafamostat. **b** Reaction progress curves of residual dasTMPRSS2 peptidase activity after 10 s pre-incubation with the indicated concentrations of inhibitor in the presence of 100 µM Boc-QAR-AMC substrate. Plateaus within progress curves demonstrate time-dependent acylation resulting in complete inhibition of activity. **c** $IC_{50}$ plots of camostat (left) and nafamostat (right) at increasing lengths of inhibitor pre-incubation, enabling calculation of kinact and Ki via Equations 2 and 3 (Methods). **d** HexaPro S protein digestion with 300 nM dasTMPRSS2 after pre-incubation with 100:1 to 0.00001:1 inhibitor:enzyme for 15 minutes. 4 µg protein were loaded per well and visualized with Coomassie blue. Kinetic plots are shown as mean ± s.e.m., *n* = 3 biological replicates and SDS-PAGE results were consistent with n=3 replicates.

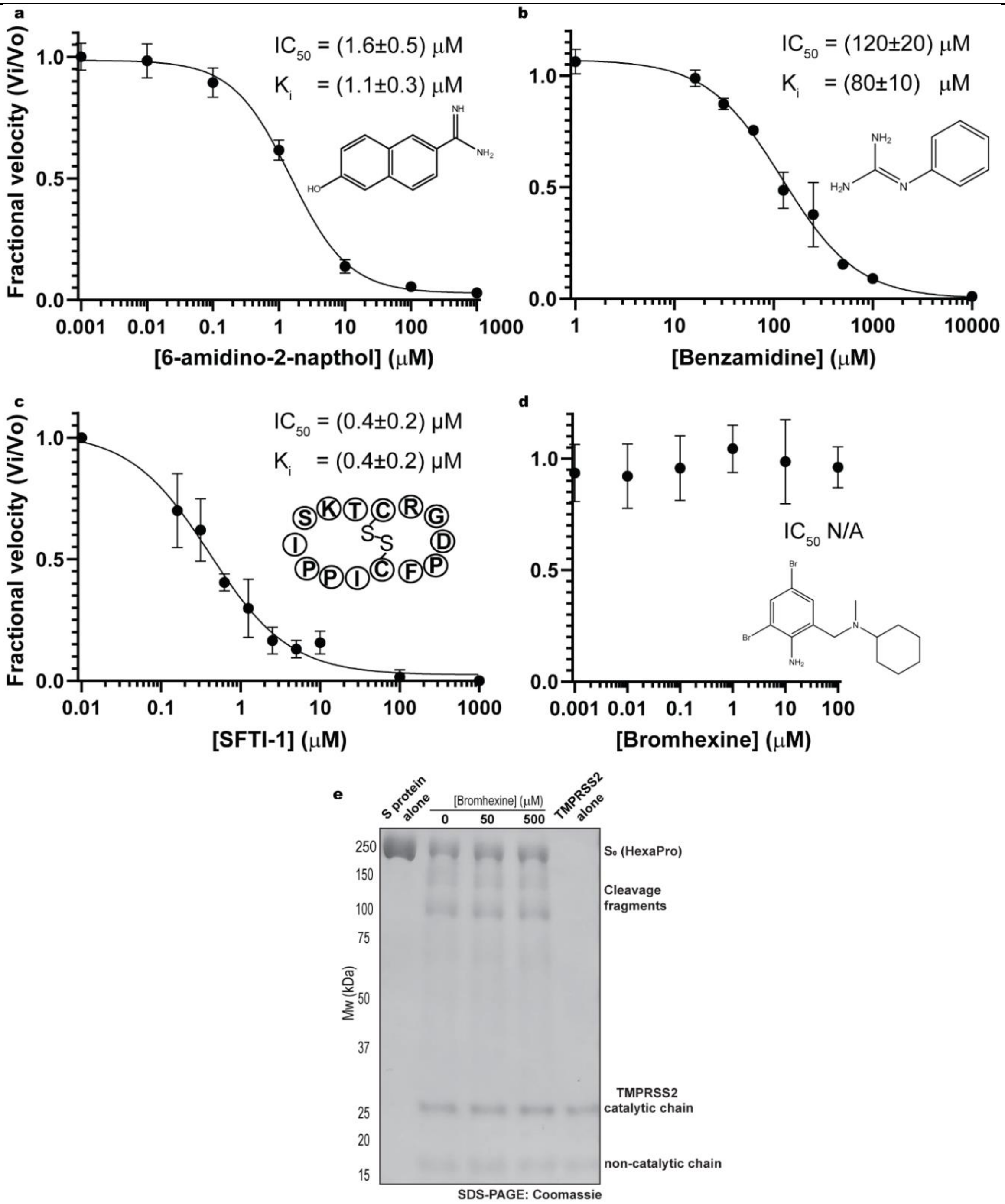

**Extended Data Fig. 7 Small molecule and peptide-based inhibitors disable dasTMPRSS2 activity unlike the clinical candidate bromhexine. a** $IC_{50}$ plot of the nafamostat leaving group, 6-amidino-2-napthol, in the presence of 100 µM Boc-QAR-AMC **b** $IC_{50}$ plot of the bicyclic peptide inhibitor, sunflower trypsin inhibitor-1 (SFTI-1) in the presence of 10 µM Boc-QAR-AMC. Synthesis and purification of this compound is available in Supplementary Information. **c** $IC_{50}$ plot of the small molecule benzamidine inhibitor in the presence of 100 µM Boc-QAR-AMC substrate. **d** IC50 plot of bromhexine in the presence of 10 µM Boc-QAR-AMC substrate showing no detectable inhibition. **e** Co-incubation of bromhexine with 300 nM TMPRSS2 results in no inhibition of proteolytic activity towards SARS-CoV-2 S protein construct HexaPro over 30 minutes. All data are shown as mean ± s.e.m., *n* = 3 or 4 biological replicates.

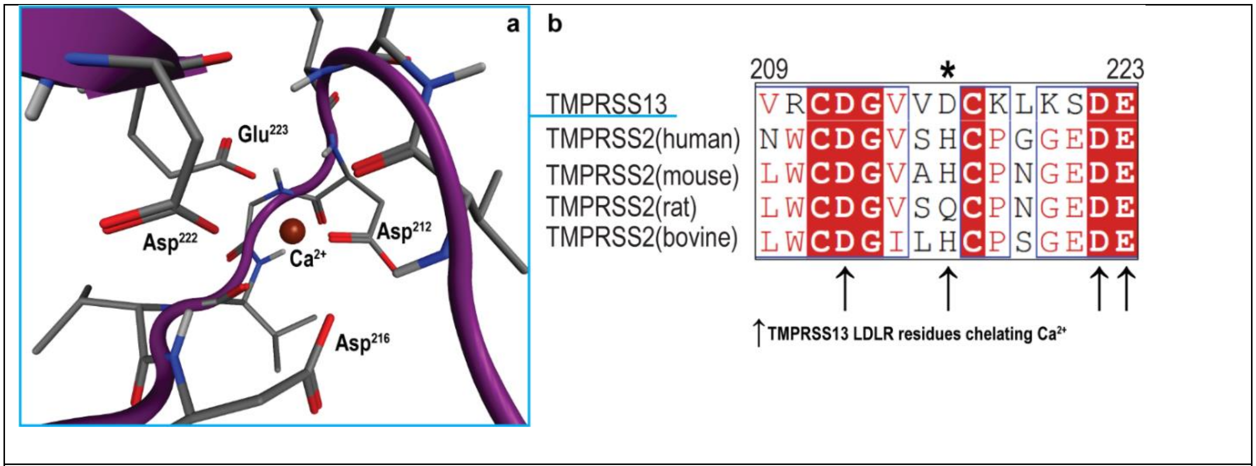

**Extended Data Fig. 8 A key calcium binding residue in TMPRSS13 is absent in TMPRSS2. a** Calcium binds within the LDLR domain of TMPRSS13 through a network of acidic residues. **b** A key calcium binding residue in TMPRSS13, Asp216 (starred), is substituted with a His or Gln residue in mammalian TMPRSS2 orthologs.
