## Supplementary Information for "Structure, activity and inhibition of human TMPRSS2, a protease implicated in SARS-CoV-2 activation"

### Detailed dasTMPRSS2 enzyme inhibition screening protocol

Commercial protease inhibitors were screened for dasTMPRSS2 inhibition in this study with the intention of developing a protocol for high throughput inhibitor screening with novel chemical libraries using nanomolar concentrations of enzyme. To minimize autolytic degradation of active enzyme, 10,000x enzyme stocks at 32 µM were prepared as 2 µL aliquots flash frozen and stored at -80 °C until immediately prior to use (Methods). Clinical protease inhibitors were dissolved to 100 mM concentrations (or the highest achievable concentration solubilized with ultrasonic bath sonication) in DMSO immediately prior to use, serving as 100x stocks to minimize the amount of DMSO added to the reaction mixture. Inhibitor assays were performed in 96-well black microplates at 200 µL volume in our study but consistent inhibitor potencies were obtained in 384-well black microplates at 50 µL volume (Data not shown). A substrate concentration of 100 µM Boc-QAR-AMC (prepared similarly to inhibitors as a 100x stock at 10 mM) was selected as it was sufficiently below the $K_{\mathrm{M}}$ to detect inhibition by weak competitive binders while also providing high fluorescent turnover across the first 60 seconds of assay initiation. A substrate and inhibitor Master Mix was prepared for each of 7 log-fold concentrations of inhibitor and compared to an uninhibited, DMSO control sample for each assay (Supplementary Table 1). Substrate + Inhibitor Master Mixes were prepared in quadruplet and dispensed to wells prior to thawing dasTMPRSS2 enzyme aliquots. Then, an Enzyme Master Mix was prepared and transferred to wells via multichannel pipette and kinetic reads were immediately performed with the fastest available read settings at ambient machine temperature (24 °C). Fluorescent values were corrected by subtracting fluorescence at assay initiation (t = 0), then initial reaction velocities ($V_0$) were determined by measuring the linear slope across the first 60 seconds of the progress curve. Normalized, fractional velocities ($V_i/V_0$) were determined by dividing inhibitor reaction velocities ($V_i$) by velocities obtained with uninhibited enzyme and plotted to obtain $\mathrm{IC}_{50}$ values.

**Supplementary Table 1**. Enzyme assay components for rapid dasTMPRSS2 inhibitor screening in 96 well or 384 well microplate formats.

| **Substrate + Inhibitor Master Mix (75% of well)** | | |
|---|---|---|
| Vol (µL; 96 well) | Vol (µL; 384 well) | **Component** |
| 2 | 0.5 | 100x substrate (or DMSO) |
| 2 | 0.5 | 100x inhibitor (or DMSO) |
| 146 | 36.5 | Assay Buffer |
| **150** | **37.5** | **Total Vol (µL)** |
| **Enzyme Master Mix (25% of well)** | | |
| Vol (µL; 96 well) | Vol (µL; 384 well) | **Component** |
| 2 | 0.5 | 100x enzyme |
| 48 | 12 | Assay Buffer |
| **50** | **12.5** | **Total Vol (µL)** |

Using this assay format (or by further downscaling to nanolitre volumes appropriate for 1536-well fluorescent plates), we anticipate libraries of compounds could be rapidly screened at 1 µM or 10 µM concentrations and compared to 1 µM nafamostat positive controls and uninhibited enzyme negative controls in order to identify novel inhibitory compounds blocking dasTMPRSS2 activity.

**SFTI-1 synthesis and purification**

Reagents and solvents were purchased from commercial sources and used without further purification, unless otherwise stated. High performance liquid chromatography (HPLC) was performed on 1) an Agilent 1260 infinity system equipped with a model 1200 quaternary pump and a model 1200 UV absorbance detector or 2) an Agilent 1260 Infinity II Preparative System equipped with a model 1260 Infinity II preparative binary pump, a model 1260 Infinity variable wavelength detector (set at 220 nm), and a 1290 Infinity II preparative open-bed fraction collector. The HPLC column used for analysis was a Phenomenex Luna $C_{18}$ semi-preparative column (5 μ, 250 × 10 mm). The HPLC column used for synthesis was a preparative column (Gemini, NX-C18, 5 μm, 110 Å, 50x30 mm) purchased from Phenomenex. Mass analyses were performed using a Waters 2695 Separation module and Waters-Micromass ZQ mass spectrometer system.

Peptides were synthesized on a Liberty Blue automated microwave peptide synthesis (CEM Corporations). Fmoc-Gly-OH was loaded on a 2-chlorotrityl resin (AdvancedChemTech) in $CH_2Cl_2$ and 2,4,6-trimethylpyridine at room temperature. The resin was capped with a solution of 10/5/85 MeOH/DIEA/$CH_2Cl_2$ for 10 minutes. Fmoc groups were removed using 20% Piperidine in DMF at 50ºC for 10 mins. Using HATU/DIEA chemistry (5 eq each in DMF) at 50ºC for 10 mins per cycle, Fmoc-Asp(OBno)-OH, Fmoc-Pro-OH, Fmoc-Phe-OH, Fmoc-Cys(ACM)-OH, Fmoc-Ile-OH, Fmoc-Pro-OH, Fmoc-Pro-OH, Fmoc-Ile-OH, Fmoc-Ser(tBu)-OH, Fmoc-Lys(Boc)-OH, Fmoc-Thr(tBu)-OH, Fmoc-Cys(ACM)-OH and Fmoc-Arg(Pbf)-OH were loaded sequentially. Fmoc-Arg(Pbf)-OH and Fmoc-amino acids coupled after Pro were coupled using two cycles. The peptide was cleaved from the resin using 20% HFIP in $CH_2Cl_2$ with no subsequent purification. The peptide was cyclized in solution using 3 eq of PyAOP and 6 eq of DIEA in DMF using microwave heating at a concentration of 1 mg/mL at 90ºC for 15 mins. The reaction was concentrated *in vacuo*. Next, the cyclized peptide was treated with 95/2.5/2.5 TFA/TIS/$H_2O$ and stirred for 3 hours. The crude peptide mixture was concentrated and added to a solution of cold diethyl ether. The suspension was centrifuged at 2500 RPM for 7 mins, the supernatant diethyl ether was discarded, and the solids were diluted into water, frozen and lyophilized to yield a white powder. The reaction was purified using preparative HPLC, eluted with 16-36% acetonitrile in water with 0.1% TFA over 20 mins at a flow rate of 30 mL/min. The retention time was 11.23 min and the collected fractions were frozen and lyophilized. Next, the collected white powder was dissolved in 300 μL of 50% AcOH in $H_2O$ and 30 μL of 1 M HCl. 10 μL of 0.1 M $I_2$ in AcOH was added to the reaction mixture and stirred for 5 hours at room temperature, while monitored using semi-preparative HPLC. After confirmation of starting material consumption, the reaction was added to a 5 mL solution of 0.1 M NaOAc and purified using preparative HPLC, eluted with 21-41% acetonitrile in water with 0.1% TFA over 20 mins at a flow rate of 30 mL/min. The retention time was 10.13 min and the collected fractions were frozen and lyophilized to yield a white powder. ESI-MS: calculated $[M+2H]^{2+}$ for $C_{67}H_{106}N_{18}O_{18}S_2$ 757.37, found 757.94.
